# Spatio–temporal model reduces species misidentification bias of spawning eggs in stock assessment of spotted mackerel in the western North Pacific

**DOI:** 10.1101/2020.04.22.056671

**Authors:** Yuki Kanamori, Shota Nishijima, Hiroshi Okamura, Ryuji Yukami, Mikio Watai, Akinori Takasuka

## Abstract

Species identification based on morphological characteristics includes species misidentification, leading to estimation bias of population size. The eggs of spotted mackerel *Scomber australicus* and chub mackerel *S. japonicus* in the western North Pacific has been identified based on egg diameter. Recent density of spotted mackerel was considerably high despite its low stock biomass. A possibility of this phenomenon is due to overestimation because the difference in egg diameter has become ambiguous between two species. However, we cannot test this possibility using DNA analysis because the eggs are fixed with formalin. Here, we estimated the index of egg density of spotted mackerel using a spatio–temporal model that incorporates the effect of egg density of chub mackerel on the catchability of spotted mackerel, using 15 years data of spawning eggs. We then examined how retrospective biases in estimated stock abundance were reduced when using the index from the model. The index estimated from the model decreased temporal fluctuation and showed smooth patterns. Especially, the recent index was considerably revised down rather than the nominal index. Additionally, the retrospective bias decreased ca. half compared with the nominal index. Therefore, incorporating species misidentification bias should be an essential process for improving stock assessment.

## 1 Introduction

Species identification based on morphological characteristics in field surveys is a major method in ecology, despite the increasing use of DNA techniques in recent years. Although most surveys are conducted under the assumption that species will be identified perfectly, this is not always the case (Elphic, 2008). Species misidentification can lead to serious bias in the inference of population size, resulting in a misunderstanding of the ecological processes that drive population dynamics. Therefore, removing the bias due to species misidentification as much as possible is essential in ecology, but such bias has drawn considerably less research attention compared with detection bias (e.g., MacKenzie et al., 2002; Williams, Nichols & Conroy, 2002).

Accurate species identification of fish eggs and larvae is essential for elucidating the ecology of the early life–history of fish, including the location and timing of fish spawning, hatching, and migration (Ko et al., 2013). Such information can improve the inference and forecasting of fish population size. Morphological characteristics used for species identification have traditionally been the size and oil globules of eggs, and the body shape, pigmentation, and meristic count of larvae (e.g., Matarese & Sandknop, 1984; Ko et al., 2013). However, species identification based on these morphological characteristics leads to species misidentification because these morphological characteristics are likely to overlap among species in early life–history (e.g., Victor et al., 2009; Ko et al., 2013). For example, when we use size of eggs as a morphological measure, we often classify eggs by whether their diameters are greater than or less than a predetermined value. However, because distributions of diameter are likely to overlap among species, some eggs may be erroneously classified as different species. In addition, morphological characteristics can change during developments, so that individuals of the same species at different development stages can be misidentified as a different species (Ko et al., 2013).

Spotted mackerel *Scomber australasicus* and chub mackerel *Scomber japonicus* are small pelagic fish that are widely distributed in the western North Pacific (ca. 120 − 150*°*E, Fig. 1; Watanabe & Yatsu, 2006). These species spawn in waters near the Kuroshio Current from winter to summer (e.g., Watanabe, 1970; Watanabe et al., 1999; Watanabe & Yatsu, 2006), after which the adults and their offspring are transported to their feeding ground by the Kuroshio Current (e.g., Watanabe & Nishida, 2002). Because Nishida (2001) suggested that there was the difference in egg diameter between two species, species identification based on egg diameter has been conducted routinely since 2005; eggs smaller than 1.1 mm of diameter were classified as chub mackerel and vice versa. These eggs, which were identified according to this basis, have been used as the indices of spawning stock biomass of spotted mackerel and chub mackerel for stock assessment. However, recent egg density of spotted mackerel was considerably high although stock biomass and spawning stock biomass has been low (Yukami et al. 2019). This considerable increase of the egg density of spotted mackerel is likely the result of overestimation because the difference in egg diameter has become ambiguous according to increase of egg density of chub mackerel and the distributions of egg diameter between species have overlapped (Yukami et al., 2019). From the possibility of overestimation, it is problematic to use a yearly trend simply estimated from the egg density data as a spawning stock biomass index for stock assessment, which could lead to bias sources in stock assessment.

**Fig. 1:**
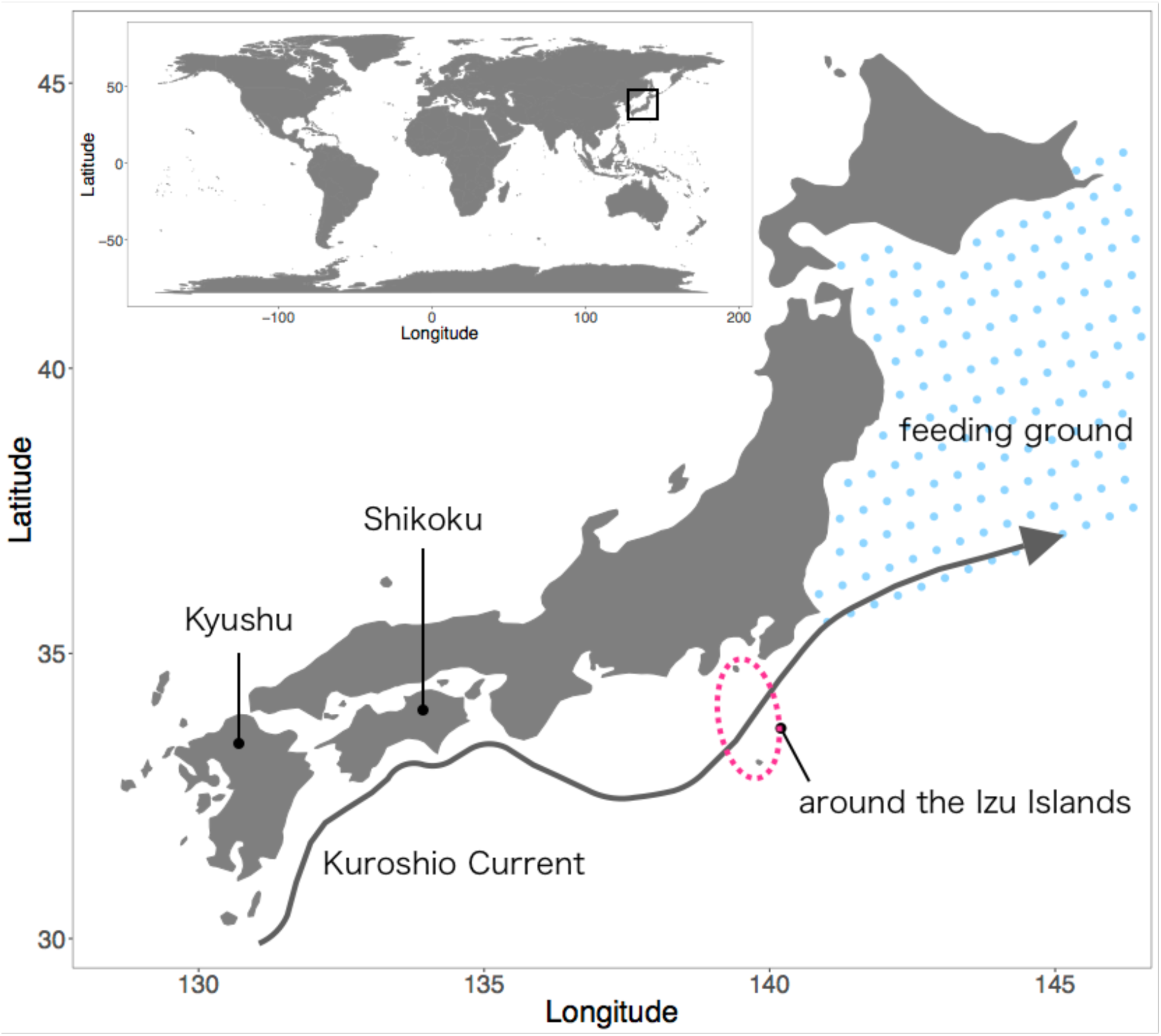
Study area. Spotted mackerel *Scomber australasicus* in the western North Pacific spawns around Kyushu, Shikoku, and the Izu Islands in Japan. Adults and their offspring are then transported to their feeding ground by the Kuroshio Current.

There are two straightforward approaches to solving this problem. The first approach is DNA analysis. However, Egg samples are fixed with formalin to preserve their morphological characteristics and this results in DNA fragmentation and protein cross-linking, which makes DNA extraction difficult or impossible (e.g., Goelz et al., 1985; Impraim et al., 1987). The second approach is to use a mixture distribution of the eggs of chub mackerel and spotted mackerel which contains the temporal changes in egg diameters of two species. However, this is difficult on a practical level because complexly intertwined factors such as spawning times within a given year, age, water temperature, and body condition affect egg diameter, and these may be difficult to obtain by field surveys alone (e.g., spawning times). As another solution, we modelled the species identification error by linking the catchability of egg density of spotted mackerel to the egg density of chub mackerel, because the recent increase of chub mackerel abundance may give rise to the identification error for spotted mackerel egg. That is, an unexpected increase of the egg density of spotted mackerel is virtually replaced by the increase of catchability of the spotted mackerel eggs.

In this paper, we demonstrate a pretty good handling of identification error by using the state-of-the-art spatio–temporal standardization method (Thorson 2019). Our new method substantially reduced the bias that would have been caused by species misidentification of spawning eggs between chub mackerel and spotted mackerel and led to considerable improvement in the stock assessment of spotted mackerel in the western North Pacific. To quantify the effect of species misidentification, we estimated the indices of egg density of spotted mackerel with/without incorporation of the effect of the egg density of chub mackerel on the catchability of spotted mackerel, using 15 years data of spawning eggs. We then examined how retrospective biases of three measurements of stock abundance (total number of individuals, total stock biomass, and spawning stock biomass; SSB) changed when we used the estimated indices for a stock assessment model. We tested the hypothesis that the retrospective bias should be lower in the spotted mackerel stock assessment with the egg–abundance index standardized by the spatio–temporal model incorporating chub mackerel egg density as a catchability covariate.

## 2 Materials and Methods

### 2.1 Data sets

#### Survey and data

The egg density data with 30^*′*^ latitude *×* 30^*′*^ longitude horizontal square resolution in the areas from 122*°*E to 150*°*E and 24*°*N to 43*°*N was used. The egg density data set was derived from monthly egg surveys off the Pacific coast of Japan from January to June, 2005–2019 (Takasuka et al., 2008a, 2019). The aim of the surveys was to monitor the egg abundance of major small pelagic fish species, including chub mackerel and spotted mackerel, so that the spatial area and survey month of the data largely covered the major spawning grounds and spawning season. While some sampling locations were fixed, others varied for various reasons (e.g., environmental conditions). Accordingly, the survey design changed slightly each year (Kanamori et al., 2019). Although the sampling efforts were approximately consistent year-round, the efforts tended to be more intensive during early spring; effort was highest in February and decreased gradually thereafter (Takasuka et al., 2008b).

The egg surveys were conducted by 18 prefectural experimental stations or fisheries research institutes and two national research institutes of the Japan Fisheries Research and Education Agency, following the consistent sampling designs, as a part of the stock assessment project. In the surveys, plankton nets were towed vertically from a depth of 150 m to the surface (if the depth was ¡150 m, nets were lowered to just above the bottom). This range of depths covers the vertical distributions of eggs of small pelagic fish. During the period from 2005 to 2019, the surveys used a plankton net with a mouth ring diameter of 0.45 m and a mesh size of 0.335 (partially 0.330 mm in 2015) (Takasuka et al., 2017). The samples were fixed with 5% formalin immediately after collection. In the laboratory, the samples were identified and sorted into eggs and larvae of different small pelagic species, based on the morphological characteristics (e.g., egg shape and size, number of oil globules, segmented yolk, perivitelline space ranging, yolk diameter, oil globule diameter). For the mackerel eggs, the egg diameters were measured to the nearest 0.025 mm by a micrometer for the maximum number of 100 individuals per sample (station or tow). Eggs with diameters *>*1.1 mm were identified as spotted mackerel, whereas those with diameters *leq*1.0 mm were identified as chub mackerel, according to Nishida et al. (2001). For any sample of *>*100 individuals, the proportion of the two species among the randomly selected 100 individuals was assumed to be the same for the whole sample.

### 2.2 Data analyses

#### Indices of egg density

In this study, we used the three indices of egg density; nominal, chub–, and chub+. The nominal index was the arithmetic mean of egg density for each year. The chub– index was the estimated egg density by considering sampling effects (i.e., spatio—temporal changes in survey design). The chub+ index was the estimated egg density by considering sampling effects and the effect of egg density of chub mackerel on the catchability of egg density of chub mackerel. The process for estimating chub– and the chub+ is described in the following section.

#### Estimation of the indices of egg density

To estimate the chub– and the chub+ indices of egg density by considering sampling effects (i.e., spatio–temporal changes in survey design) as well as the effect of egg density of chub mackerel on the catchability of egg density of chub mackerel, we used the multivariate vector autoregressive spatio-temporal (VAST) model (Thorson & Barnett, 2017), which accounts for spatio-temporal changes in survey design, survey effort, and observation rates and can accurately estimate relative local densities at high resolution by standardizing sampling designs (Thorson & Barnett, 2017; Thorson, 2019). The model includes two potential components because it is designed to support delta-models: (i) the encounter probability *p*_*i*_ for each sample *i* and (ii) the expected egg density *d*_*i*_ for each sample *i* when spawning occurs (i.e., egg density is not zero). The encounter probability *p*_*i*_ and the expected egg density *d*_*i*_ are, respectively, approximated using a logit-linked linear predictor and a log-linked linear predictor as follows (Thorson & Barnett, 2017):

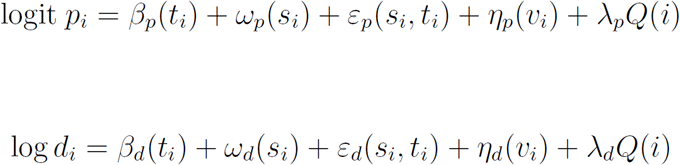

where *β*(*t*_*i*_) is the intercept for year *t*, and *ω*(*s*_*i*_) and *ε*(*s*_*i*_, *t*_*i*_) are the spatial and spatio–temporal random effects for year *t* and location *s*, respectively. *η*(*v*_*i*_) is overdispersion random effect of factor *v*_*i*_ which is the interaction of year and month. *λ* is the effect of the chatchability covariate *Q*(*i*), where *Q*(*i*) = log(chub mackerel egg density(*s*_*i*_) + 0.1). That is, this term considers the effect of species misidentification between chub mackerel and spotted mackerel; as mentioned earlier, we suspected overestimation of egg density of spotted mackerel because the difference in egg diameter has become ambiguous according to increase of egg density of chub mackerel and the distributions of egg diameter between species have overlapped (Yukami et al., 2019). The subscripts for each term on the right side, *p* and *d*, represent the encounter probability and the expected egg density, respectively.

The probability density function of *ω*(*·*) is a multivariate normal distribution MVN(0, **R**), where the variance–covariance matrix **R** is a Matérn correlation function. The probability density function of *ε*(*s*_*i*_, *t*_*i*_) is

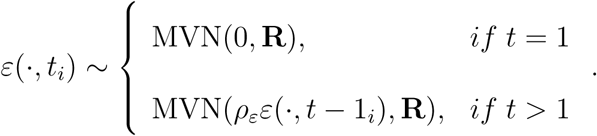

Here, *ρ*_*ε*_ = 0 because we assumed that the year was independent. Therefore, the probability density function of *η*(*v*_*i*_) is *η*(*v*_*i*_) ∼ N(0, 1).

For computational reasons, the spatio-temporal variation *ε*_*p*_(*s*_*i*_, *t*_*i*_) was approximated as being piecewise constant at a fine spatial scale. We used a k-means algorithm to identify 200 locations (termed “knots”) to minimize the total distance between the location of sampling data (Thorson et al., 2015) using R-INLA software (Lindgren, 2012). The number of knots was increased to the greatest extent possible, and similar results were obtained for low knots (= 100; Akaike information criterion [AIC] = 6773.01) and high knots (= 200; AIC = 6676.25).

Parameters in the VAST model were estimated using the VAST package (Thorson et al., 2015,2016a) in R 3.6.1 (R Development Core Team, 2019). Bias-correction for random effects (Thorson and Kristensen, 2016) was applied when estimating the derived parameters. We confirmed the model diagnostics plots and found no serious problems. The relative egg density in year *t* at location *s*, 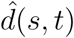 and the index of egg density in year *t*, 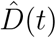, were estimated using the predicted values for random effects as follows (Thorson et al., 2017):

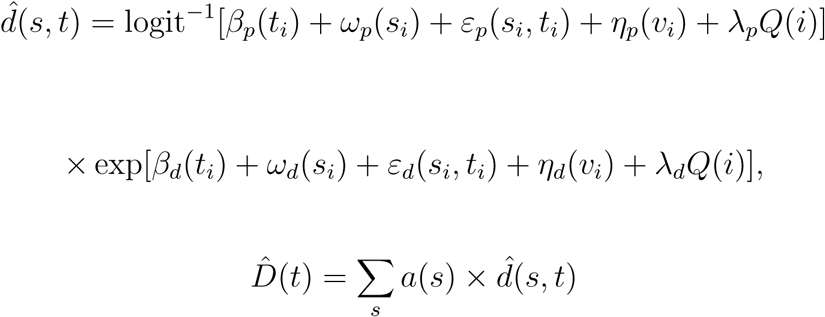

where *a*(*s*) is the area of location *s*.

#### Estimation of stock abundance

To examine the validity of the three indices (i.e., nominal index, chub-index, and chub+ index), we estimated the three measurements of stock abundance (total number of individuals, total stock biomass, and SSB) using a tuned virtual population analysis (VPA). This model is an age-based cohort analysis for estimating the historical abundance and fishing mortality rates from catch-at-age data and has been applied to spotted mackerel in Japan (Yukami et al., 2019). In addition to the three indices of egg density, we used catch-at-age, weight-at-age, maturity-at-age, the natural mortality coefficient, and a recruitment index following stock assessment in Japan (Yukami et al., 2019). The fishing mortality coefficients other than the terminal age in the terminal year were estimated under the assumption that the selectivity in the latest year was equal to the average selectivity of the prior 5 years (Ichinokawa & Okamura, 2014; Mori & Hiyama, 2014). We confirmed that this assumption did not change our results when using the prior 3 years average of selectivity as the selectivity in the latest year. The fishing mortality coefficient at each age in the terminal year was estimated by a maximum likelihood method as follows:

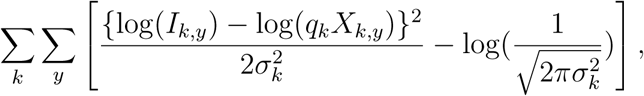

where *I*_*k,y*_ is the value of index *k* in year *y, q*_*k*_ is a proportionality constant, *X*_*k,y*_ is the abundance estimate in VPA for index *k* (i.e., recruitment, and the three indices of egg density), 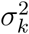 is the variance in fitting the abundance estimate to the index, and *y*_*k*_ is the first year of index *k*.

#### Retrospective analysis

Stock abundance in the terminal year estimated by VPA is notoriously inaccurate and imprecise compared with historical abundance estimates (Okamura et al. 2017). One of the most serious problems is that the stock abundance estimate in the terminal year has temporally systematic bias, i.e., retrospective bias (Hurtado-Ferro et al. 2015). Retrospective analysis is therefore a useful method for detecting such a systematic bias in stock abundance estimate in the terminal year. Dropping the most recent year’s data sequentially and then comparing the estimates from a full-year data model and removed data model reveals presence or absence of systematic bias (Mohn 1999). Herein, we conduct a retrospective analysis to evaluate the relative goodness of the three indices of egg density.

To examine improvements in estimations of the three measurements of stock abundance when using the estimated indices of egg density from VAST with considering the chub mackerel’s effect, we performed a retrospective analysis by sequentially removing the five most recent years of data from the full data set. Retrospective analysis is usually used in stock assessment models such as VPA to examine the reliability and predictability of stock assessments (e.g., Mohn, 1999; Hashimoto et al., 2018). We calculated Mohn’s rho to estimate the biases of the indices of egg density as follows (Mohn, 1999):

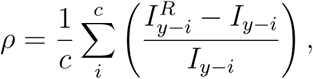

where *I*_*y*−*i*_ is the value of the year *y* − *i* estimate using the full data and 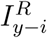 is the estimate using the data up to year *y* − *i. c* is the maximum number of removed years (i.e., *c* = 5). A positive *ρ* means that the estimate in the terminal year tends to be positively biased on average, and vice versa. Moreover, a *ρ* close to 0 means no serious retrospective bias and greatly improved estimation of the stock abundance.

## 3 Results

### Temporal trend in the indices of egg density

When comparing the standardized indices to the nominal index, the standardized indices reduced temporal fluctuation and showed smooth patterns (Fig. 2). Whereas the nominal index increased substantially in 2018, the standardized indices were revised downward to a considerable degree. Moreover, the standardized indices of some years, such as 2008, 2009, and 2012, were revised upward.

**Fig. 2:**
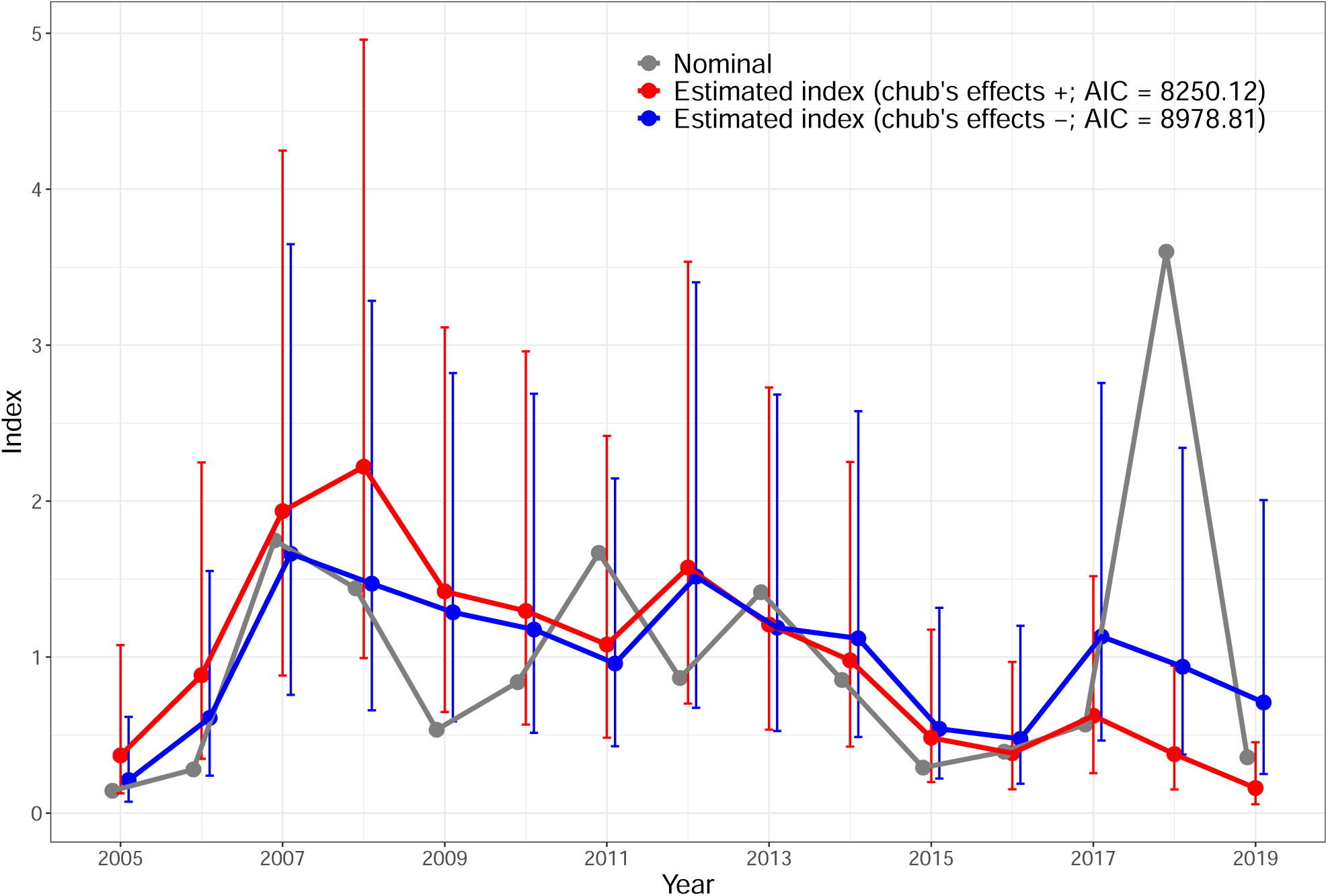
Temporal trend of the indices of egg density. The grey line represents the scaled nominal index, the blue line represents the estimated index without the chub mackerel effect, and the red line represents the estimated index with chub mackerel effect. Vertical bars are 95% confidence intervals of the estimated indices.

The model with the effect of chub mackerel’s egg density on the catchability of spotted mackerel was more suitable model rather than the model without the effect of chub mackerel’s, which based on AIC criteria (chub+, AIC = 8250.12; chub–, AIC = 8978.81). The coefficient of the effect of chub mackerel’s on the catchability of spotted mackerel, *λ*, represents a positive effect (*λ* = 0.17). The estimated index with the chub mackerel’s effect reached a peak in 2008 and then gradually decreased. The value of this index in 2019 was the lowest since 2005 (Fig. 2).

### Spatial distribution of the relative egg density

The relative egg density with the effect of chub mackerel was high off the coast of Kyushu, Shikoku, and the Izu Islands (Fig. 3). In addition, the relative egg density was slightly high off the coast of the Tohoku region. These tendencies were consistent during the study period. There was no area where the relative egg density clearly increased or decreased during this study period.

**Fig. 3:**
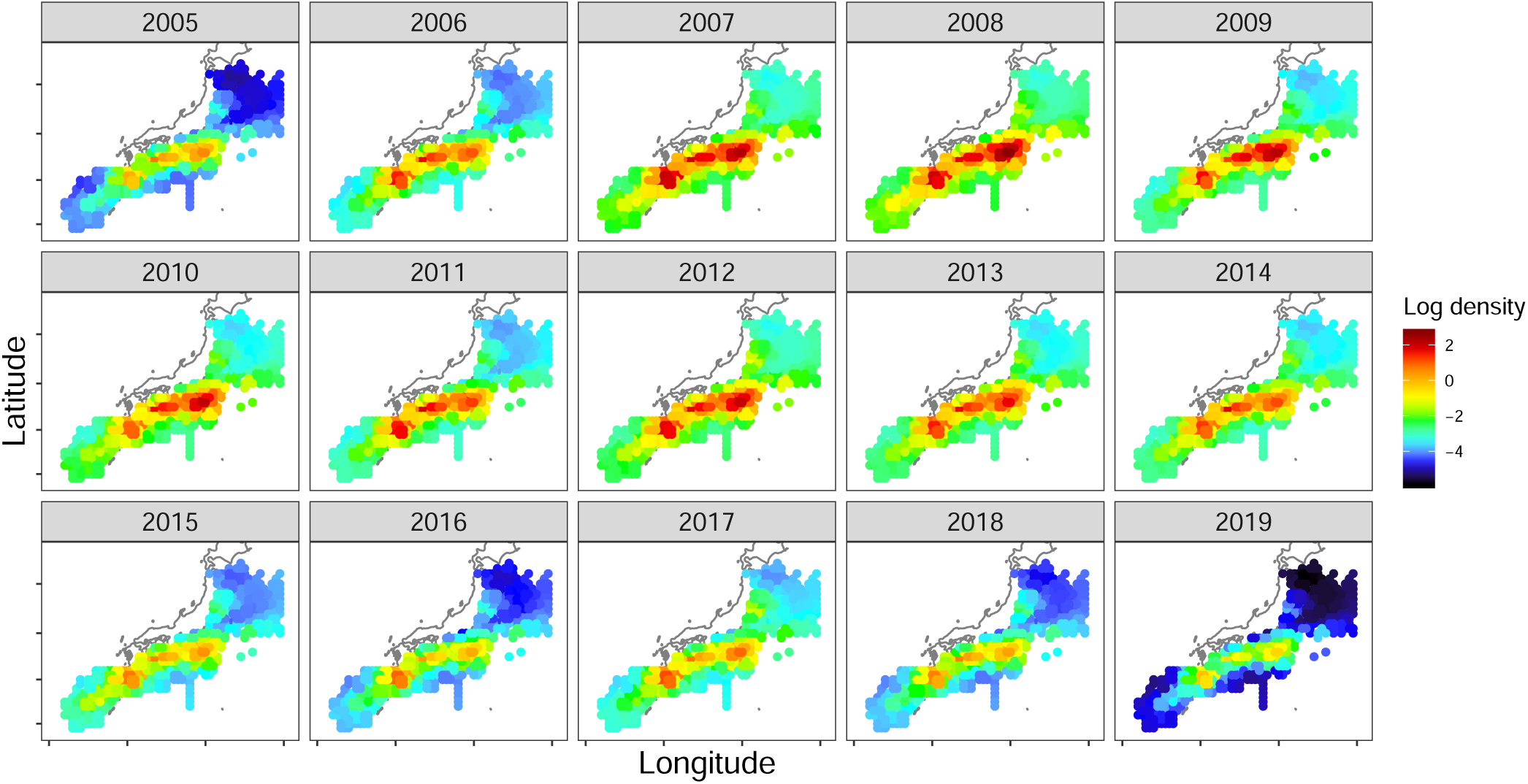
Temporal changes in the spatial distribution of relative egg density, which estimated by using the model with chub mackerel effect.

### Retrospective analysis

Recent estimated values of stock abundance (i.e., total numbers of individuals, total biomass, and SSB) were distinct depending on what indices were used, whereas the directions of retrospective bias were sometimes not, depending on the indices used (Fig. 4). In all the three measurements of stock abundance, the recent estimated values were higher when using the nominal and estimated index without the chub mackerel’s effect rather compared with using the estimated index with the chub mackerel’s effect. The directions of retrospective bias were always positive and were independent if the indices used.

**Fig. 4:**
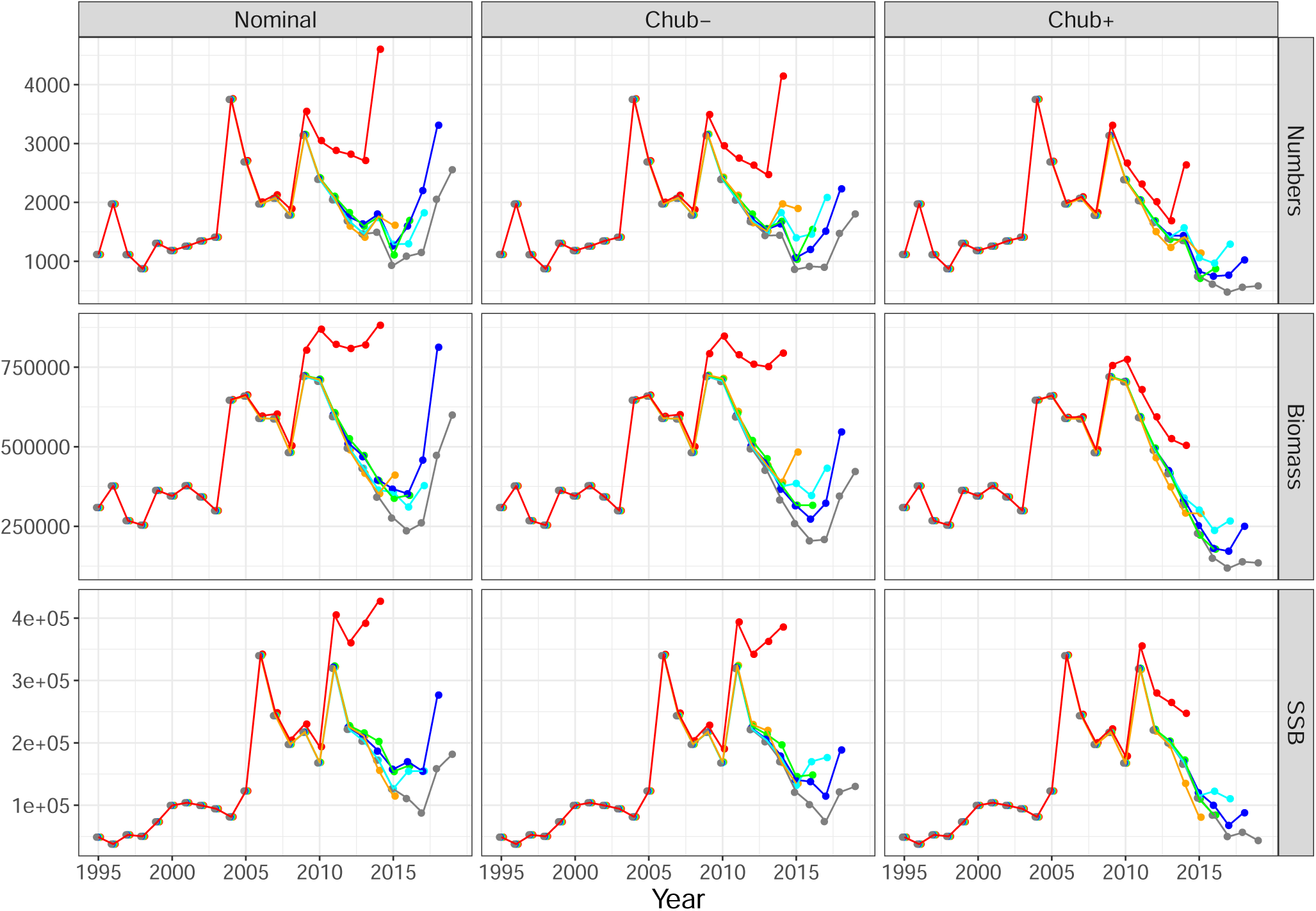
Retrospective patterns of total numbers of individuals, total biomass, and spawning stock biomass (SSB).

In all the three measurements of stock abundance (i.e., total numbers of individuals, total biomass, and SSB), retrospective biases were clearly improved when using the estimated index with the chub mackerel’s effect (Table 1). Mohn’s rho, which represents the magnitude and direction of retrospective bias, had similar values between when using nominal index as when using the estimated index without the chub mackerel’s effect (Table 1). In contrast, Mohn’s rho decreased when using the estimated index with the chub mackerel effect. The directions of the retrospective bias did not change depending on the indices used because the values of Mohn’s rho were always positive.

**Table 1:** Mohn’s rho for each index of total numbers of individuals, total biomass, and spawning stock biomass (SSB).

| Index | Mohn's rho |  |  |
| --- | --- | --- | --- |
|  | Numbers | Biomass | SSB |
| Nominal | 0.47 | 0.45 | 0.44 |
| Chub – | 0.51 | 0.48 | 0.45 |
| Chub + | 0.28 | 0.24 | 0.19 |

## 4 Discussion

We modelled the species identification error by linking the catchability of egg density of spotted mackerel to the egg density of chub mackerel. We found that the model incorporating the effect of the egg density of chub mackerel was the better model, based on AIC (Fig. 2). In addition, the model showed a positive effect of the egg density of chub mackerel on the catchability of spotted mackerel. These results suggest the necessity of incorporating the effect of the egg density of chub mackerel when standardizing the egg density of spotted mackerel.

Methods that reduce the bias in species misidentification are needed for accurate stock assessment because inaccurate estimates of stock size may lead to incorrect management decisions and endanger exploited populations in the long term (Marko et al., 2004; Garcia-Vazque et al., 2012). The retrospective biases in all the three measurements of stock abundance were clearly improved when using the estimated index with the chub mackerel effect; the magnitude of the retrospective biases decreased by about half compared with when the other indices were used (Fig. 4 and Table 1). These results suggest that our new method is effective for reducing the bias in species misidentification and greatly improves the stock estimation especially of pelagic eggs, which have less significant differences in shape and size for species identification. Species samples for preserving morphological characteristics are usually fixed with formalin because of some advantages such as small shrinks of tissue and low cost than with ethanol. However, it is difficult to extract DNA from formalin-fixed samples due to DNA fragmentation and protein cross-linking (e.g., Goelz et al., 1985; Impraim et al., 1987), and so DNA analysis of these samples for species identification is difficult, if not impossible. Accordingly, species samples which were collected prior to development of DNA techniques cannot used for DNA analysis. In contrast, our new method requires only the geographic locations and “prior-” information, such as the species name (which can be based on some morphological characteristics), to be able to use various data such as the survey data of eggs and larvae collected in the ICES area. Thus, our method should be of great benefit to fisheries science.

Our results can play an important role on actual management of spotted mackerel. The stock status and management of this species is the focus of much attention in Japan because this species is one of the nine TAC (total allowable catch) species, whose catches are strictly managed according to output control. In fact, a new harvest control rule based on maximum sustainable yield (MSY) was implemented in 2020 (Yukami et al. 2020). The stock abundance of spotted mackerel has been decreasing in recent years, and positive retrospective bias caused overestimation of abundance in the terminal year in previous stock assessment using the nominal index of spawning egg (Yukami et al. 2019). This indicates that the allowable biological catch (ABC) was also overestimated, and this may have led to overfishing. The present study found that the retrospective bias was considerably mitigated by incorporating the effect of mixing of chub mackerel’s eggs on spotted mackerel’s egg and, thus, would contribute to the derivation of ABC at an adequate level. Although the current status is overfishing and overfished (Yukami et al. 2020), it is expected that the Pacific stock of spotted mackerel will show a recovery to a level that produces MSY, using our assessment method and the new Harvest Control Rules.

The geographic location of spawning grounds did not change in spotted mackerel (Fig. 3), whereas the geographical location of spawning grounds has been shifted northward in chub mackerel (Kanamori et al., 2019). This difference in change of spawning ground between the two species may make it more difficult to perform species identification based on egg diameter because the diameter of marine fish eggs generally increases in higher latitudes (Llanos–Rivera & Castro, 2004). In other words, the egg diameter of chub mackerel may increase as their spawning ground shifts northward, making it closer in size to that of the spotted mackerel. This suggests that rising sea temperatures associated with climate change may affect not only spatio–temporal patterns of organisms, such as phenology and spatial distribution, but also an estimation of population abundance.

Although detailed information on spawning grounds is necessary for understanding of the fluctuations in recruitment as well as a basis for stock management, prior data on the spotted mackerel has not been reliable. For example, some studies have reported that the waters around the Izu Islands may not be a suitable spawning ground for spotted mackerel because few eggs have been observed (Yukami et al. 2019). In contrast, it is possible that the spotted mackerel spawns around the Izu Islands because the estimated hatch day and the spatial distribution of spotted mackerel at the Kuroshio–Oyashio transition area were similar to those of chub mackerel, which spawns around mainly the Izu Islands (Takahashi et al., 2010). The present study showed that the relative egg density, which was estimated using the better model, was equally high off the coast of Kyushu, Shikoku, and the Izu Islands (Fig. 3), providing direct evidence that the waters around the Izu Islands are also a major spawning ground of spotted mackerel. One reason that spotted mackerel spawn in the waters around the Izu Islands is that spotted mackerel are not sensitive to rising water temperatures because they are generally distributed farther south than chub mackerel (Mitani et al., 2002). Indeed, although both spotted mackerel and chub mackerel spawn at the same time around the Izu Islands (Tanoue et al., 1960; Hanai & Meguro, 1997), the reproductive phenology of chub mackerel has changed due to rising sea surface temperatures associated with climate change; chub mackerel have been migrating to their feeding ground earlier and spawning father northward since 2000 (Kanamori et al., 2019).

Understanding migration patterns is necessary for conducting stock assessments (Crossin et al., 2017). It has been assumed that spotted mackerel changes their spawning ground with age; spotted mackerel migrates from around the Izu Islands to the Kuroshio–Oyashio transition area to feed before spawning at 2 years of age (Nishida et al., 2000; Kawabata et al. 2008). Adults that have spawned gradually migrate westward, using the spawning grounds off the coast of Kyushu and Shikoku (Hanai, 1999; Nashida et al., 2006). Although the number of recruits was substantially high in 2004 and 2009 (Yukami et al., 2019), we did not find a tendency toward increased the relative egg density around the Izu Islands in 2006 and in 2011 or the other spawning grounds after 2007 and 2012 (Fig. 3). One explanation for this is the possibility that the migration range of spotted mackerel is narrower than we assumed. Previous studies have reported that spotted mackerel has retention around the Izu Islands and off the coast of Shikoku (Hanai, 1999; Nashida et al., 2006). Another explanation is that part of a strong year may remain in another area due to the expansion of spatial distribution resulting from an increased number of recruitments. For example, Kawabata et al. (2008) reported that the 2004 year class migrated for feeding and overwintering until at least 3 years old over the Emperor Seamounts (around 165 − 170*°*E and 30 − 55*°*N). Testing these hypotheses will be the subject of future research and should improve our understanding of the migratory patterns of the spotted mackerel, which in turn should improve stock assessment and management.

## Conclusion

This study showed that the indices of egg density of spotted mackerel, which were standardized using a spatio–temporal model, reduced temporal fluctuation and showed smooth patterns. In particular, the standardized indices in 2018 were revised downward to a considerable degree compared with the nominal index. The model incorporating the effect of chub mackerel egg dens ity on the catchability of spotted mackerel (i.e., the model incorporating species misidentification bias) was the better model according to the AIC criteria. In addition, the retrospective bias decreased by about half when using the egg density index from the better model. These results suggest that incorporating species misidentification bias should be an essential process in improving stock assessment.

## Acknowledgments

This research was financiallysupportted by the grants from the Japan Society for the Promotion of Science (JSPS) (19K15905, 20392904).

